# Dissociable Default Mode Network Connectivity Patterns Underlie Distinct Symptoms in Psychosis Risk

**DOI:** 10.1101/2024.10.25.620271

**Authors:** Chelsea C. Ajunwa, Jiahe Zhang, Guusje Collin, Lihua Xu, Matcheri S. Keshavan, Yingying Tang, Tianhong Zhang, Huijun Li, Martha E. Shenton, William S. Stone, Jijun Wang, Margaret Niznikiewicz, Susan Whitfield-Gabrieli

**Affiliations:** Department of Psychology, Northeastern University, Boston, MA; McGovern Institute for Brain Research, Massachusetts Institute of Technology, Cambridge, MA; Radboudumc, Department of Psychiatry, Nijmegen, the Netherlands; Donders Institute for Brain, Cognition and Behaviour, Nijmegen, the Netherlands; Shanghai Key Laboratory of Psychotic Disorders, Shanghai Mental Health Center, Shanghai Jiao Tong University School of Medicine, Shanghai, China; Department of Psychiatry, Beth Israel Deaconess Medical Center, Harvard Medical School, Boston, MA; Department of Psychology, Florida A&M University, Tallahassee, FL; Psychiatry Neuroimaging Laboratory, Department of Psychiatry, Brigham and Women’s Hospital, Harvard Medical School, Boston, MA; Research and Development, VA Boston Healthcare System, Brockton Division, Brockton, MA; Department of Radiology, Brigham and Women’s Hospital, Harvard Medical School, Boston, MA; Department of Psychiatry, VA Boston Healthcare System, Brockton Division, Brockton, MA

## Abstract

The Clinical High Risk (CHR) stage of psychosis is characterized by subthreshold symptoms of schizophrenia including negative symptoms, dysphoric mood, and functional deterioration. Hyperconnectivity of the default-mode network (DMN) has been observed in early schizophrenia, but the extent to which hyperconnectivity is present in CHR, and the extent to which such hyperconnectivity may underlie transdiagnostic symptoms, is not clear. As part of the Shanghai At-Risk for Psychosis (SHARP) program, resting-state fMRI data were collected from 251 young adults (158 CHR and 93 controls, M = 18.72, SD = 4.68, 129 male). We examined functional connectivity of the DMN by performing a whole-brain seed-to-voxel analysis with the MPFC as the seed. Symptom severity across a number of dimensions, including negative symptoms, positive symptoms, and affective symptoms were assessed. Compared to controls, CHRs exhibited significantly greater functional connectivity (p < 0.001 uncorrected) between the MPFC and 1) other DMN nodes including the posterior cingulate cortex (PCC), and 2) auditory cortices (superior and middle temporal gyri, STG/MTG). Furthermore, these two patterns of hyperconnectivity were differentially associated with distinct symptom clusters. Within CHR, MPFC-PCC connectivity was significantly correlated with anxiety (r= 0.23, p=0.006), while MPFC-STG/MTG connectivity was significantly correlated with negative symptom severity (r=0.26, p=0.001). Secondary analyses using item-level symptom scores confirmed a similar dissociation. These results demonstrate that two dissociable patterns of DMN hyperconnectivity found in the CHR stage may underlie distinct dimensions of symptomatology.

## Introduction

Individuals who are diagnosed with schizophrenia typically suffer from a combination of affective, cognitive, and psychomotor symptoms.^1^ Schizophrenia onset is marked by the first episode of psychosis, which typically occurs in adolescence or young adulthood.^2^ Prior to onset, many individuals experience a psychosis-risk, or Clinical High-Risk (CHR) syndrome, which is characterized by subthreshold symptoms of schizophrenia.^3^ These syndromes may include negative symptoms (e.g., avolition, alogia, anhedonia)^4,5^, dysphoric mood,^6^ impaired self representation,^7^ cognitive decline,^8^ and functional deterioration in major life roles,^6^ among others.

Recent advances in neuroimaging have enabled the study of functional connectivity, i.e., the statistical interdependence of blood-oxygen-level-dependent (BOLD) signals between brain regions. Functional connectivity has been characterized in a variety of clinical populations, including individuals with schizophrenia.^9^ Large-scale disruptions in functional connectivity have been reported in chronic schizophrenia,^10–12^ early-stage psychosis,^10,13–16^, and CHR.^17,18^

More specifically, previous studies have reported hyperactivity and hyperconnectivity within the default mode network (DMN),^19–22^ an intrinsic brain network which includes regions such as the medial prefrontal cortex (MPFC), posterior cingulate cortex (PCC), and angular gyri (AG). The DMN is active at rest and during internally focused and self-referential tasks.^23,24^ Furthermore, stronger DMN functional connectivity is associated with worse psychotic symptoms in patients and in individuals with familial risk.^22^ In CHR individuals, there is evidence of increased DMN connectivity,^25,26^ although no association with symptom severity has been reported.

The auditory cortices, including the superior and middle temporal gyri (STG/MTG), also exhibit altered neurobiology in schizophrenia, including differences in brain volume and cortical thickness,^27–31^ cerebral blood flow,^32,33^ white matter microstructure,^34,35^ activation,^36,37^ and functional connectivity^27,38^ compared to healthy controls. A recent meta-analysis found that STG and MTG are among the main regions with structural alterations in gray matter volume and white matter integrity in schizophrenia patients with persistent negative symptoms.^39^ Evidence also suggests that connectivity between the STG/MTG and DMN regions such as the MPFC is associated with negative symptoms in schizophrenia.^40^ In addition to being associated with negative symptoms, DMN-STG/MTG connectivity may also be associated with another aspect of schizophrenia symptomatology, auditory verbal hallucinations (AVH). The “resting state hypothesis” of AVH was developed based on observations of elevated resting state activity in both the auditory cortex and anterior cortical midline DMN regions such as the MPFC. It posits that MPFC-STG hyperconnectivity underlies misattribution of internally generated phenomena as external.^41^ However, evidence of MPFC-STG/MTG hyperconnectivity has not yet been reported in clinical high risk individuals or those with schizophrenia.

In the present study, we examined rsfMRI data in CHR and healthy control (HC) participants from the Shanghai At-Risk for Psychosis (SHARP) program. We hypothesized that compared to HCs, CHR participants would exhibit hyperconnectivity within the DMN as well as between the DMN and auditory regions. Within CHR participants, we tested associations between DMN functional connectivity and summary scores of psychotic and affective symptoms assessed via the Structured Interview for Psychosis-Risk Syndromes (SIPS), Hamilton Anxiety Rating Scale (HAM-A), and Hamilton Depression Rating Scale (HAM-D). We further explored associations between DMN functional connectivity and item-level symptom scores.

## Methods and Materials

### Participants

This study involved 158 CHR participants and 93 healthy controls (HCs) matched on age, sex, and education. Most were naïve to psychotropic medication at baseline clinical assessment (95%) and neuroimaging (80%). The CHR participants were help-seeking individuals who were recruited at the Shanghai Mental Health Center (SMHC) for possible inclusion in the Shanghai At Risk for Psychosis Program (SHARP). Healthy control participants were recruited through advertisements. SHARP is a collaborative, international, NIMH-funded research effort^18,42–55,55–72^ between the SMHC and Harvard Medical School at Beth Israel Deaconess Medication Center (BIDMC) and Brigham and Women’s Hospital, the Massachusetts Institute of Technology, and Northeastern University. The program was approved by the Institutional Review Boards at both BIDMC and the SMHC. All subjects or their legal guardians provided written informed consent. Participants under 18 also provided assent. **Table 1** provides basic demographic and clinical information for the current sample, similar to previous publications that utilized SHARP resting state data.^18,44^

**Table 1.** Demographic and clinical characteristics.

|  | **Controls (N=93)** | **CHR (N=158)** | **Statistics** |
| --- | --- | --- | --- |
| Age, mean (sd) [range] | 18.7 (4.6) [12 - 35] | 18.8 (4.9) [13 - 34] | $t=-0.43$, $p=0.671$ |
| Sex, male/female | 49 / 44 | 80/78 | $\chi^2=0.03$, $p=0.854$ |
| Education in years, mean (sd) [range] | 10.8 (2.3) [6 - 17] | 10.5 (2.8) [4 - 19] | $t=0.88$, $p=0.381$ |
| IQ, mean (sd) [range] | 104.2 (11.1) [75 - 133] | 98.9 (12.8) [52 - 128] | $t=3.24$, $p=0.001$ |
| Framewise displacement, mean (sd) [range] | 0.1 (0.1) [0.0 - 0.4] | 0.1 (0.0) [0.0 - 0.3] | $t=1.25$, $p=0.214$ |
| Psychotropic medication |  |  |  |
| At inclusion, N (%) |  | 7 (4.4) |  |
| At baseline MRI, N (%) |  | 29 (18.4) |  |
| Antipsychotics, N (%) |  | 24 (15.2) |  |
| Antidepressants, N (%) |  | 7 (4.4) |  |
| Other, N (%) |  | 2 (1.2) |  |
| Baseline SIPS scores |  |  |  |
| Positive, mean (sd) [range] | 0.4 (0.8) [0-4] | 10.1 (3.6) [0 - 21] | $t=-32.25$, $p<.001$ |
| Negative, mean (sd) [range] | 0.3 (0.8) [0-4] | 11.7 (6.1) [1 - 27] | $t=-23.05$, $p<.001$ |
| Disorganized, mean (sd) [range] | 0.3 (0.5) [0-2] | 6.6 (3.2) [1 - 19] | $t=-24.01$, $p<.001$ |
| General, mean (sd) [range] | 0.6 (1.0) [0-4] | 9.1 (3.2) [1 - 17] | $t=-30.79$, $p<.001$ |
| Total, mean (sd) [range] | 1.6 (1.9) [0-9] | 37.4 (10.8) [13 - 79] | $t=-40.46$, $p<.001$ |
| Hamilton Anxiety Rating Scale Score |  | 8.9 (4.3) [1 - 22] |  |
| Hamilton Depression Rating Scale Score |  | 14.3 (7.3) [3 - 46] |  |

### Clinical and cognitive assessment

Prodromal symptoms were assessed using a validated Chinese version of the Structured Interview for Psychosis-Risk Syndromes (SIPS)^61,73^, including 19 items grouped into 4 subscales (SIPS-P: positive symptoms; SIPS-N: negative symptoms; SIPS-D: disorganization symptoms; SIPS-G: general symptoms). A Chinese version of the Scale of Psychosis Symptoms (SOPS) was derived from the SIPS to determine CHR staging. A score on the global assessment of functioning (GAF) scale was also generated based on the SIPS.^74^ SIPS and GAF scores were collected at a baseline assessment and at a one year follow-up visit. The MINI International Neuropsychiatric Interview (MINI) was used to identify previous and current psychotic disorders. Current symptoms of anxiety and depression were assessed using the Hamilton Anxiety Rating Scale (HAM-A) and the Hamilton Depression Rating Scale (HAM-D), respectively. Overall cognitive ability (i.e., IQ) was estimated with the Wechsler Abbreviated Scale of Intelligence (WASI).^75^

### Image acquisition

Magnetic resonance imaging (MRI) scans were acquired using a 3T Siemens MR B17 (Verio) system with a 32-channel head coil. A T1-weighted anatomical MRI scan was taken (MP-RAGE; TR = 2300 ms, TE = 2.96 ms, FA = 9°, FOV = 256 mm, voxel size 1 × 1 × 1 mm^3^, 192 contiguous sagittal slices, duration 9′14′′). A resting-state scan was also taken (149 functional volumes; TR = 2500 ms, TE = 30 ms, FA = 90°, FOV = 224 mm, voxel size 3.5 × 3.5 × 3.5 mm^3^, 37 contiguous axial slices, duration 6′19′′).

### Image preprocessing

We preprocessed scans using the CONN toolbox.^76^ In brief, preprocessing steps included segmentation of gray and white matter tissue, realignment, slice-timing correction, normalization to Montreal Neurological Institute (MNI) space, and smoothing with a 6mm FWHM Gaussian filter (see details in Collin et al 2020). Potential spurious correlations in rsfMRI time-series were assessed using the Artifact Detection Tool (ART; http://www.nitrc.org/projects/artifact_detect). Outliers, defined as volumes with head displacement in the x, y, or z greater than 1mm relative to the previous frame or a mean global intensity greater than 3 standard deviations from the mean intensity for the whole rsfMRI scan, were removed from the data via linear regression. Motion correction was performed using 3 rotational, 3 translational, and 1 composite motion parameters. The resulting time-series were corrected for artifactual covariates and signals within white matter (3 principal components) and cerebrospinal fluid (3 principal components). They were also band-pass filtered (0.008 Hz-0.09 Hz). We calculated averaged framewise displacement across all timepoints for each participant.^77^

### Between-group functional connectivity analysis

We first created an *a priori* MPFC seed (**Figure 1A**): an 8mm sphere centered at MNI coordinates (-3, 44, -2).^78^ Using the CONN toolbox, we computed Pearson’s correlation coefficients between the average time course within the seed and the time courses of all other voxels in the brain. We converted Pearson’s *r values* to *z*-scores via Fisher’s transform. We then performed a two-sample t-test (CHR vs. HC) to identify voxels whose functional connectivity to the MPFC was significantly different between groups. A voxel threshold of *p*<.001 (uncorrected) and a cluster threshold of *p*<0.05 (controlling for false discovery rate, or FDR^79^) were used. For further analysis, we extracted connectivity strength (Fisher transformed *z*-scores) between the MPFC seed and three clusters, i.e., the posterior cingulate cortex (PCC) as well as left and right STG/MTG (averaged to derive a single MPFC-STG/MTG connectivity measure). To determine if there was an effect of medication status on MPFC-PCC or MPFC-STG/MTG connectivity, we performed a t-test comparing medicated subjects to non-medicated subjects.

**Figure 1.**
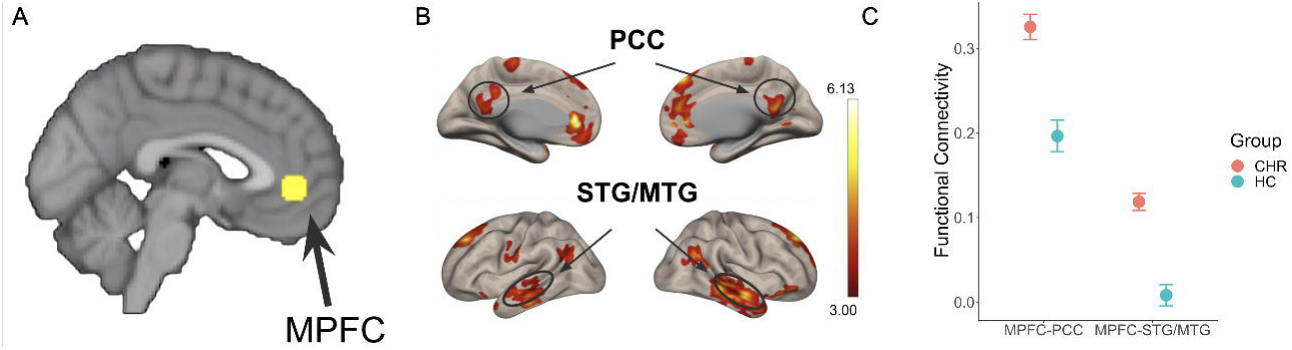
MPFC hyperconnectivity in CHR. **A)** We used an 8mm spherical seed in the MPFC.^78^ **B)** A whole-brain search revealed hyperconnectivity between the MPFC seed and several regions, including auditory cortices and the posterior cingulate cortex (*p* <.001, uncorrected). **C)** Group-level summary statistics for MPFC-STG/MTG and MPFC-PCC functional connectivity are displayed in magenta for CHR and in cyan for HC. Error bars indicate 1 standard error of the mean.

Next, we performed the above procedure again using other key nodes of the DMN as seeds. A PCC seed was defined *a priori* as an 8mm sphere centered at MNI coordinates (-3, -39, 39)^78^ (**Figure S1B**). An angular gyri seed was defined *a priori* as an 8mm sphere centered at MNI coordinates (-47, -70, 37; 52, -71, 38)^78^ and averaged across left and right hemispheres (**Figure S1C**). We examined the functional connectivity between these seeds and the rest of the brain in order to compare their connectivity profiles to that of the MPFC.

### Identification of connectivity-symptom relationships

To test brain-behavior relationships (in R v4.2.2.^80,81^), we ran linear regressions with each relevant clinical assessment (SIPS-P subscale, SIPS-N subscale, GAF, HAM-A, and HAM-D) as the outcome variable and functional connectivity (i.e., MPFC-PCC and MPFC-STG/MTG connectivity) as the predictor variable and performed FDR correction^79^ to account for multiple comparisons. (While SIPS-D and SIPS-G were not our primary measures of interest, we tested their associations with MPFC-PCC and MPFC-STG/MTG connectivity as well.)

The relationships between MPFC-PCC and MPFC-STG/MTG connectivity and symptom data from follow-up visits conducted one year post-baseline were also examined. SIPS-P and SIPS-N at follow-up were used as the primary psychotic symptom scales while SIPS-G2, the dysphoric mood item of the general symptoms subscale, was used as a measure of affective symptom severity (measures of HAM-A and HAM-D were not obtained at follow-up). GAF at follow-up was also examined.

### Secondary analyses of connectivity-symptom relationships

To further explore the associations between functional connectivity and symptoms, we ran two secondary analyses using all 19 SIPS items, GAF, HAM-A, and HAM-D, i.e., 22 symptom scales total. First, we ran 22 separate regressions with the symptom scales as outcome and performed FDR correction to account for multiple comparisons. For all regression analyses, we also ran sensitivity analyses controlling for age, sex and framewise displacement.

Second, we conducted a principal component analysis (PCA) in R to reduce dimensionality within the symptom scales in a data-driven manner. For each of the two PCs that accounted for the most variance in the symptom data, we summed the product of the symptom scores and the corresponding component loadings to derive a composite score for each participant. Correlations between the resulting composite scores and MPFC functional connectivity were identified via linear regression.

## Results

### Increased functional connectivity in CHR vs. HC

Compared to HC, CHR exhibited significantly greater functional connectivity (*p* < .001, uncorrected) between the MPFC seed and several regions (**Figure 1B**), including bilateral auditory cortices (middle temporal gyrus and superior temporal gyrus; left hemisphere peak: -52, -24, -20; right hemisphere peak: 56, -6, -22) and the PCC (peak: -2, -58, 6) as the two largest clusters (**Figure 1C**). Eight smaller clusters also exhibited hyperconnectivity to the MPFC seed, including additional DMN regions such as bilateral angular gyri, superior frontal gyri and left cerebellum Crus II (**Table S1**). Similar results were observed when we controlled for age, sex, and framewise displacement (**Figure S2)**. In medicated subjects compared to non-medicated subjects, there was no difference (*p* > .05) in MPFC-PCC connectivity or MPFC-STG/MTG connectivity (**Figure S3**). CHR also exhibited greater functional connectivity (*p* < .001, uncorrected) between the PCC and AG seeds and several regions (**Figure S1**). Notably, in CHR the PCC and AG were not hyperconnected to the STG/MTG region.

### Associations between functional connectivity and clinical measures

Within CHR participants, we found dissociable relationships between MPFC hyperconnectivity and symptom severity (**Figure 2**). Specifically, controlling for FDR (*q_FDR_* < .05), stronger MPFC-PCC connectivity was significantly associated with higher HAM-A scores (*r* = 0.23, *p* = 0.006), and not with the SIPS-N subscale (*r* = 0.15, *p* = 0.056; **Figure 2A**). Greater MPFC-STG/MTG connectivity was significantly (*q_FDR_* < .05) associated with higher total SIPS-N score (*r* = 0.26, *p* = 0.001), but not with HAM-A (*r* = 0.09, *p* = 0.286; **Figure 2B**). Neither MPFC-PCC nor MPFC-STG/MTG connectivity was significantly associated with HAM-D, SIPS-P or GAF (*q_FDR_* < .05).

**Figure 2.**
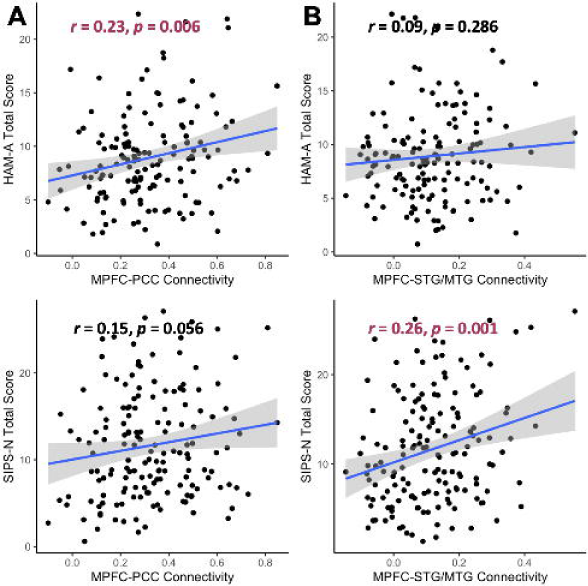
Dissociable relationships between MPFC connectivity and symptom severity in CHR. MPFC connectivity measures were defined based on CHR>HC comparison and independently of symptom measures. **A)** Individual differences in MPFC-PCC connectivity were significantly associated with variation in HAM-A scores and marginally so in SIPS-N scores. **B)** Individual differences in MPFC-STG/MTG connectivity were associated with variation in SIPS-N score, but not in HAM-A scores.

Our tests also revealed a significant association between baseline MPFC-PCC connectivity and change in SIPS-G2 between baseline and follow-up (*r* = -0.22, *p* = 0.010; **Figure S4**). There were also trend-level associations between MPFC-PCC connectivity and 1) GAF change (*r* = 0.15, *p* = 0.078) and 2) SIPS-N change (*r* = -0.16, *p* = 0.057).

Secondary analyses on SIPS items revealed that, after FDR correction (*q_FDR_* < 0.05), MPFC-PCC connectivity was significantly associated with two general symptom items (G2, Dysphoric Mood and G4, Impaired Tolerance to Normal Stress) and MPFC-STG/MTG connectivity was significantly associated with two negative symptom items (N5, Ideational Richness; N2, Avolition) (**Table S2**). See supplementary materials for a full correlation matrix for all connectivity and clinical variables (including all SIPS subscales and individual items, GAF, HAM-A, and HAM-D; **Figure S5**).

A secondary PCA revealed two PCs that together explained 69.4% of variance (PC1: 46.4%; PC2: 23.0%) in scores on the 22 symptom scales (**Figure S6A**). PC1 and PC2 were characterized by strong loadings for negative symptoms and affective symptoms (including both anxiety and depression symptoms) respectively (**Figure S6B**). Stronger MPFC-PCC connectivity was significantly associated with higher PC2 scores (*r* = 0.27*, p <* 0.001) but not with PC1 scores (*r* = -0.02*, p =* 0.821; **Figure S7A**). Stronger MPFC-STG/MTG connectivity was significantly associated with PC1 scores (*r* = -0.20*, p =* 0.016) but not with PC2 scores (*r* = 0.11*, p =* 0.106; **Figure S7B**). PC1 and PC2 variable loadings can be found in **Table S3**.

Sensitivity analyses revealed that similar dissociations were found after controlling for age, sex, and motion (**Table S2**).

## Discussion

This study examined the functional connectivity changes that occur during the clinical high-risk stage of psychosis, thereby providing insight into the mechanisms underlying the development of the disorder. Our investigation of CHR functional connectivity patterns revealed that, compared to HCs, CHR individuals exhibited hyperconnectivity 1) within the DMN and 2) between the MPFC and STG/MTG. Within the CHR group, within-DMN hyperconnectivity was preferentially associated with affective symptoms (i.e., anxiety as measured by HAM-A). MPFC-STG/MTG hyperconnectivity, on the other hand, was preferentially associated with negative symptoms (i.e., negative symptoms as measured by SIPS) and decreased global functioning.

This dissociation was further confirmed by secondary analyses of the SIPS items (**Figure 3**). Affective symptoms (i.e., dysphoric mood) were found to be preferentially associated with MPFC-PCC connectivity, while negative symptoms (i.e., avolition, ideational richness/alogia) were preferentially associated with MPFC-STG/MTG connectivity. Further, a principal component analysis identified dimensions of symptomatology within the sample. The first principal component seemed to capture variability within the negative symptom dimension of the SIPS and was associated with MPFC-STG/MTG connectivity. The second principal component seemed to capture variability within the affective dimension of CHR symptomatology and was associated with MPFC-PCC connectivity. Prior investigations into the dimensional structure of schizophrenia symptomatology have identified separate negative and affective symptom dimensions.^82–85^ Our findings suggest that, in CHR, these two dimensions may be neurobiologically distinct, involving disparate pathophysiological mechanisms.

**Figure 3.**
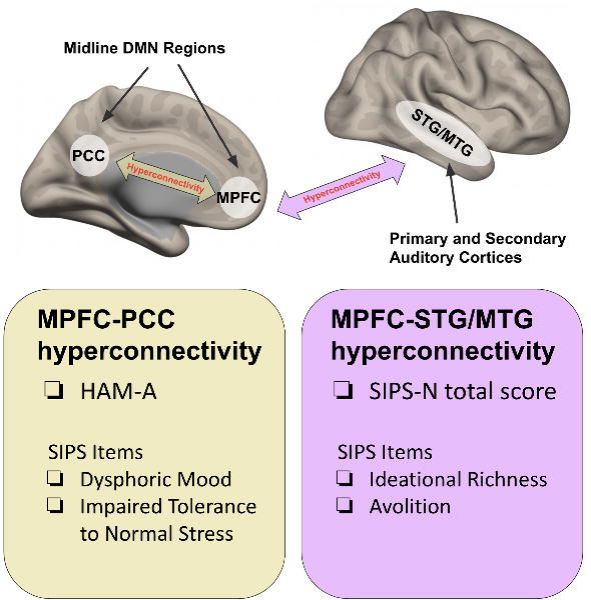
Hyperconnectivity in relation to SIPS items. MPFC-PCC hyperconnectivity in CHR was associated with HAM-A, Dysphoric Mood and Impaired Tolerance to Normal Stress. MPFC-STG/MTG hyperconnectivity in CHR was associated with SIPS-N total score, GAF, and the Ideational Richness and Avolition SIPS items.

Hyperconnectivity within the DMN has been previously reported in patients and relatives of people with schizophrenia,^20–22,86^ although the opposite, hypoconnectivity, has been observed as well.^87–89^ Importantly, within-DMN hyperconnectivity is associated with greater symptom severity^22^ and predicts worse outcome^90^ in early stage psychosis. Past studies have also reported DMN hyperconnectivity in the CHR stage of psychosis.^25,26^ The present study replicates these findings. Furthermore, our study reveals an association between baseline within-DMN hyperconnectivity and baseline affective symptoms, such as anxiety, dysphoric mood and impaired tolerance to normal stress. Within-DMN connectivity at baseline was also associated with improvement in dysphoric mood between baseline assessment and follow-up. The direction of this relationship is consistent with the observation that individuals with higher baseline pathology (i.e., more room for improvement) show more reductions in symptoms at a followup time point.^91,92,93^

Beyond schizophrenia, within-DMN hyperconnectivity was also observed in major depressive disorder,^94–98^ chronic pain,^99^ autism,^100^ and post-traumatic stress disorder.^101^ Furthermore, several studies have identified associations between DMN connectivity and affective symptom severity in these populations.^94–96,99^ More specifically, rumination, a form of repetitive negative thinking defined by a focus on one’s own negative mood, is associated with DMN hyperconnectivity in both healthy^102^ and depressed^95,98,103–105^ populations. Rumination is a transdiagnostic feature of several psychiatric illnesses including major depressive disorder, generalized anxiety disorder, post-traumatic stress disorder, substance abuse disorders, eating disorders, and insomnia.^106,107^ These findings suggest that within-DMN hyperconnectivity may be a transdiagnostic phenomenon underlying rumination across a range of psychiatric illnesses, including schizophrenia. This is consistent with evidence that some CHR individuals are later diagnosed with disorders other than schizophrenia, such as major depressive disorder and generalized anxiety disorder.^108,109^

Thus, in addition to contributing evidence of a transdiagnostic nature of DMN hyperconnectivity, the current study also provides motivation for interventions aimed at reducing MPFC-PCC hyperconnectivity and thereby ameliorating the associated symptoms across disorders. Previous studies have identified several non-invasive strategies for down-regulating the DMN, including exercise,^110^ meditation,^111^ cognitive behavioral therapy,^112^ antidepressant medications,^113^ transcranial magnetic stimulation,^114^ and real-time fMRI neurofeedback.^115^

To our knowledge, the current study is the first to report DMN-STG/MTG hyperconnectivity and further link it to negative symptoms in CHR individuals. A previous study found a similar association between fronto-temporal connectivity and negative symptoms in schizophrenia patients, although no group difference was observed between patients and HCs.^40^. The mechanism linking DMN-STG/MTG hyperconnectivity to negative symptoms has not been fully elucidated. However, there is evidence that auditory processing deficits may underlie cognitive impairment,^116,117^ which is associated with negative symptoms in schizophrenia.^118–120^ This framework linking impaired processing of auditory information with negative symptoms^121^ may be extended to the CHR stage given our current finding. At the same time, the MPFC, together with the dorsal subsystem of the DMN,^122^ is implicated in mentalizing. The pathophysiology of the DMN is also linked to poor social cognition in schizophrenia.^123^ It is noteworthy that ideational richness, the SIPS negative symptom item most associated with MPFC-STG/MTG connectivity in our CHR group, measures inability to follow conversations and articulate thoughts.^73^ Hyperconnectivity between the MPFC and STG/MTG may therefore contribute to negative symptoms via combined impairments in auditory processing and social cognition.^40^

Our novel hyperconnectivity findings in CHR may also provide support for the ‘resting state hypothesis’ of AVH, which posits that elevated resting state activity in the auditory cortex is a result of abnormal interactions between the auditory cortex and anterior DMN midline regions at rest.^41^ This hypothesis is based on evidence of hyperactivity in the STG and MTG at rest, hyperactivity of the anterior DMN at rest, and decreased stimulus-induced auditory cortex activity in patients with schizophrenia.^22,32,37,124,125^ Consistent with the resting state hypothesis, after real-time neurofeedback training aimed at reducing DMN activity, patients with schizophrenia exhibited reduced MPFC-STG connectivity associated with reduced AVH.^126^ With our current study, we report the first evidence that an abnormal interaction between MPFC and STG is present in the prodromal stage of the disorder. Furthermore, we show that, within the DMN, hyperconnectivity with the STG/MTG is specific to the anterior cortical midline portion (MPFC) as hypothesized. However, it is noteworthy that we did not find any association between MPFC-STG/MTG connectivity and the SIPS item for hallucinations (P4: Perceptual abnormalities/hallucinations). This may be due to subthreshold AVH presence in CHR. The onset of negative symptoms, cognitive deficits and resultant functional impairment often precede the development of positive symptoms and the development of psychosis.^5,6,127,128^ The broad nature of the SIPS P4 measure, which captures perceptual abnormalities and hallucinations across all sensory modalities, may also explain the lack of association.

The present study has identified distinct and dissociable patterns of DMN hyperconnectivity as potential biomarkers for affective and negative dimensions of prodromal symptom development. Future research should focus on tracking DMN hyperconnectivity and its associated symptoms over time, as a longitudinal design coupled with deep phenotyping of within-individual changes can inform on conversion vs. remission trajectories, enable mechanistic mapping of observed brain-behavior links (e.g., MPFC-STG/MTG hyperconnectivity, negative symptoms and cognitive deficits), and suggest optimal time points for intervention.

## Author Contributions

CCA and JZ contributed equally as co-first authors. CCA and JZ had full access to all the data in the study and take responsibility for the integrity of the data and the accuracy of the data analysis.

## References

1. Tandon R, Nasrallah HA, Keshavan MS. Schizophrenia, “just the facts” 4. Clinical features and conceptualization. Schizophr Res. 2009;110(1):1–23. doi:10.1016/j.schres.2009.03.005

2. Cannon TD. How Schizophrenia Develops: Cognitive and Brain Mechanisms Underlying Onset of Psychosis. Trends Cogn Sci. 2015;19(12):744–756. doi:10.1016/j.tics.2015.09.009

3. Cornblatt B, Obuchowski M, Roberts S, Pollack S, Erlenmeyer–Kimling L. Cognitive and behavioral precursors of schizophrenia. Dev Psychopathol. 1999;11(3):487–508. doi:10.1017/S0954579499002175

4. Kirkpatrick B, Fenton WS, Carpenter WT Jr, Marder SR. The NIMH-MATRICS Consensus Statement on Negative Symptoms. Schizophr Bull. 2006;32(2):214–219. doi:10.1093/schbul/sbj053

5. Piskulic D, Addington J, Cadenhead KS, et al. Negative symptoms in individuals at clinical high risk of psychosis. Psychiatry Res. 2012;196(2):220–224. doi:10.1016/j.psychres.2012.02.018

6. Yung AR, McGorry PD. The Prodromal Phase of First-episode Psychosis: Past and Current Conceptualizations. Schizophr Bull. 1996;22(2):353–370. doi:10.1093/schbul/22.2.353

7. Nelson B, Thompson A, Yung AR. Basic Self-Disturbance Predicts Psychosis Onset in the Ultra High Risk for Psychosis “Prodromal” Population. Schizophr Bull. 2012;38(6):1277–1287. doi:10.1093/schbul/sbs007

8. Catalan A, Salazar de Pablo G, Aymerich C, et al. Neurocognitive Functioning in Individuals at Clinical High Risk for Psychosis: A Systematic Review and Meta-analysis. JAMA Psychiatry. 2021;78(8):859–867. doi:10.1001/jamapsychiatry.2021.1290

9. Zhang J, Kucyi A, Raya J, et al. What have we really learned from functional connectivity in clinical populations? NeuroImage. 2021;242:118466. doi:10.1016/j.neuroimage.2021.118466

10. Li T, Wang Q, Zhang J, et al. Brain-Wide Analysis of Functional Connectivity in First-Episode and Chronic Stages of Schizophrenia. Schizophr Bull. 2017;43(2):436–448. doi:10.1093/schbul/sbw099

11. Lynall ME, Bassett DS, Kerwin R, et al. Functional Connectivity and Brain Networks in Schizophrenia. J Neurosci. 2010;30(28):9477–9487. doi:10.1523/JNEUROSCI.0333-10.2010

12. Venkataraman A, Whitford TJ, Westin CF, Golland P, Kubicki M. Whole brain resting state functional connectivity abnormalities in schizophrenia. Schizophr Res. 2012;139(1-3):7–12. doi:10.1016/j.schres.2012.04.021

13. Anticevic A, Hu X, Xiao Y, et al. Early-Course Unmedicated Schizophrenia Patients Exhibit Elevated Prefrontal Connectivity Associated with Longitudinal Change. J Neurosci. 2015;35(1):267–286. doi:10.1523/JNEUROSCI.2310-14.2015

14. Hummer T, Yung M, Goni J, et al. Functional network connectivity in early-stage schizophrenia. Schizophr Res. 2020;218:107–115. doi:10.1016/j.schres.2020.01.023

15. Kim A, Ha M, Kim T, et al. Triple-Network Dysconnectivity in Patients With First-Episode Psychosis and Individuals at Clinical High Risk for Psychosis. Psychiatry Investig. 2022;19(12):1037–1045. doi:10.30773/pi.2022.0091

16. Kim M, Kim T, Ha M, Oh H, Moon SY, Kwon JS. Large-Scale Thalamocortical Triple Network Dysconnectivities in Patients With First-Episode Psychosis and Individuals at Risk for Psychosis. Schizophr Bull. Published online December 1, 2022:sbac174. doi:10.1093/schbul/sbac174

17. Cao H, Chung Y, McEwen SC, et al. Progressive reconfiguration of resting-state brain networks as psychosis develops: Preliminary results from the North American Prodrome Longitudinal Study (NAPLS) consortium. Schizophr Res. 2020;226:30–37. doi:10.1016/j.schres.2019.01.017

18. Collin G, Seidman LJ, Keshavan MS, et al. Functional connectome organization predicts conversion to psychosis in clinical high-risk youth from the SHARP program. Mol Psychiatry. 2020;25(10):2431–2440. doi:10.1038/s41380-018-0288-x

19. Liu H, Kaneko Y, Ouyang X, et al. Schizophrenic Patients and Their Unaffected Siblings Share Increased Resting-State Connectivity in the Task-Negative Network but Not Its Anticorrelated Task-Positive Network. Schizophr Bull. 2012;38(2):285–294. doi:10.1093/schbul/sbq074

20. Sasabayashi D, Takahashi T, Takayanagi Y, et al. Resting state hyperconnectivity of the default mode network in schizophrenia and clinical high-risk state for psychosis. Cereb Cortex. 2023;33(13):8456–8464. doi:10.1093/cercor/bhad131

21. Tang J, Liao Y, Song M, et al. Aberrant Default Mode Functional Connectivity in Early Onset Schizophrenia. PLOS ONE. 2013;8(7):e71061. doi:10.1371/journal.pone.0071061

22. Whitfield-Gabrieli S, Thermenos HW, Milanovic S, et al. Hyperactivity and hyperconnectivity of the default network in schizophrenia and in first-degree relatives of persons with schizophrenia. Proc Natl Acad Sci U S A. 2009;106(4):1279–1284. doi:10.1073/pnas.0809141106

23. Buckner RL, Andrews-Hanna JR, Schacter DL. The Brain’s Default Network. Ann N Y Acad Sci. 2008;1124(1):1–38. doi:10.1196/annals.1440.011

24. Raichle ME, MacLeod AM, Snyder AZ, Powers WJ, Gusnard DA, Shulman GL. A default mode of brain function. Proc Natl Acad Sci. 2001;98(2):676–682. doi:10.1073/pnas.98.2.676

25. Damme KSF, Pelletier-Baldelli A, Cowan HR, Orr JM, Mittal VA. Distinct and opposite profiles of connectivity during self-reference task and rest in youth at clinical high risk for psychosis. Hum Brain Mapp. 2019;40(11):3254–3264. doi:10.1002/hbm.24595

26. Shim G, Oh JS, Jung WH, et al. Altered resting-state connectivity in subjects at ultra-high risk for psychosis: an fMRI study. Behav Brain Funct BBF. 2010;6:58. doi:10.1186/1744-9081-6-58

27. Shinn AK, Baker JT, Cohen BM, Öngür D. Functional connectivity of left Heschl’s gyrus in vulnerability to auditory hallucinations in schizophrenia. Schizophr Res. 2013;143(2):260–268. doi:10.1016/j.schres.2012.11.037

28. Bandeira ID, Barouh JL, Bandeira ID, Quarantini L. Analysis of the superior temporal gyrus as a possible biomarker in schizophrenia using voxel-based morphometry of the brain magnetic resonance imaging: a comprehensive review. CNS Spectr. 2021;26(4):319–325. doi:10.1017/S1092852919001810

29. Honea R, Crow TJ, Passingham D, Mackay CE. Regional Deficits in Brain Volume in Schizophrenia: A Meta-Analysis of Voxel-Based Morphometry Studies. Am J Psychiatry. 2005;162(12):2233–2245. doi:10.1176/appi.ajp.162.12.2233

30. Köse G, Jessen K, Ebdrup BH, Nielsen MØ. Associations between cortical thickness and auditory verbal hallucinations in patients with schizophrenia: A systematic review. Psychiatry Res Neuroimaging. 2018;282:31–39. doi:10.1016/j.pscychresns.2018.10.005

31. Rajarethinam RP, DeQuardo JR, Nalepa R, Tandon R. Superior temporal gyrus in schizophrenia: a volumetric magnetic resonance imaging study. Schizophr Res. 2000;41(2):303–312. doi:10.1016/S0920-9964(99)00083-3

32. Homan P, Kindler J, Hauf M, Walther S, Hubl D, Dierks T. Repeated measurements of cerebral blood flow in the left superior temporal gyrus reveal tonic hyperactivity in patients with auditory verbal hallucinations: a possible trait marker. Front Hum Neurosci. 2013;7. doi:10.3389/fnhum.2013.00304

33. Zhuo C, Zhu J, Qin W, Qu H, Ma X, Yu C. Cerebral blood flow alterations specific to auditory verbal hallucinations in schizophrenia. Br J Psychiatry. 2017;210(3):209–215. doi:10.1192/bjp.bp.115.174961

34. Lee K, Yoshida T, Kubicki M, et al. Increased diffusivity in superior temporal gyrus in patients with schizophrenia: A Diffusion Tensor Imaging study. Schizophr Res. 2009;108(1):33–40. doi:10.1016/j.schres.2008.11.024

35. Spray A, Beer AL, Bentall RP, Sluming V, Meyer G. Microstructure of the superior temporal gyrus and hallucination proneness - a multi-compartment diffusion imaging study. NeuroImage Clin. 2018;20:1–6. doi:10.1016/j.nicl.2018.06.027

36. Allen P, Larøi F, McGuire PK, Aleman A. The hallucinating brain: A review of structural and functional neuroimaging studies of hallucinations. Neurosci Biobehav Rev. 2008;32(1):175–191. doi:10.1016/j.neubiorev.2007.07.012

37. Jardri R, Pouchet A, Pins D, Thomas P. Cortical Activations During Auditory Verbal Hallucinations in Schizophrenia: A Coordinate-Based Meta-Analysis. Am J Psychiatry. 2011;168(1):73–81. doi:10.1176/appi.ajp.2010.09101522

38. Alderson-Day B, McCarthy-Jones S, Fernyhough C. Hearing voices in the resting brain: A review of intrinsic functional connectivity research on auditory verbal hallucinations. Neurosci Biobehav Rev. 2015;55:78–87. doi:10.1016/j.neubiorev.2015.04.016

39. Zhu T, Wang Z, Zhou C, et al. Meta-analysis of structural and functional brain abnormalities in schizophrenia with persistent negative symptoms using activation likelihood estimation. Front Psychiatry. 2022;13:957685. doi:10.3389/fpsyt.2022.957685

40. Abram SV, Wisner KM, Fox JM, et al. Fronto-temporal connectivity predicts cognitive empathy deficits and experiential negative symptoms in schizophrenia. Hum Brain Mapp. 2017;38(3):1111–1124. doi:10.1002/hbm.23439

41. Northoff G, Qin P. How can the brain’s resting state activity generate hallucinations? A ‘resting state hypothesis’ of auditory verbal hallucinations. Schizophr Res. 2011;127(1):202–214. doi:10.1016/j.schres.2010.11.009

42. Anteraper SA, Guell X, Collin G, et al. Abnormal Function in Dentate Nuclei Precedes the Onset of Psychosis: A Resting-State fMRI Study in High-Risk Individuals. Schizophr Bull. 2021;47(5):1421–1430. doi:10.1093/schbul/sbab038

43. Chen Y, Wang J, Xu L, et al. Age-related changes in self-reported psychotic experiences in clinical help-seeking population: From 15 to 45 years. Early Interv Psychiatry. 2022;16(12):1359–1367. doi:10.1111/eip.13285

44. Collin G, Nieto-Castanon A, Shenton ME, et al. Brain functional connectivity data enhance prediction of clinical outcome in youth at risk for psychosis. NeuroImage Clin. 2020;26:102108. doi:10.1016/j.nicl.2019.102108

45. Cui H, Giuliano AJ, Zhang T, et al. Cognitive dysfunction in a psychotropic medication-naïve, clinical high-risk sample from the ShangHai-At-Risk-for-Psychosis (SHARP) study: Associations with clinical outcomes. Schizophr Res. 2020;226:138–146. doi:10.1016/j.schres.2020.06.018

46. Del Re EC, Stone WS, Bouix S, et al. Baseline Cortical Thickness Reductions in Clinical High Risk for Psychosis: Brain Regions Associated with Conversion to Psychosis Versus Non-Conversion as Assessed at One-Year Follow-Up in the Shanghai-At-Risk-for-Psychosis (SHARP) Study. Schizophr Bull. 2020;47(2):562–574. doi:10.1093/schbul/sbaa127

47. Li H, Yang S, Chi H, et al. Enhancing attention and memory of individuals at clinical high risk for psychosis with mHealth technology. Asian J Psychiatry. 2021;58:102587. doi:10.1016/j.ajp.2021.102587

48. Li H, Yang S, Chi H, et al. Functionality and feasibility of cognitive function training via mobile health application among youth at risk for psychosis. Explor Digit Health Technol. Published online February 27, 2024:7–19. doi:10.37349/edht.2024.00007

49. Song W, Xu L, Zhang T, et al. Peripheral transcriptome of clinical high-risk psychosis reflects symptom alteration and helps prognosis prediction. Psychiatry Clin Neurosci. 2022;76(6):268–270. doi:10.1111/pcn.13346

50. Tang Y, Pasternak O, Kubicki M, et al. Altered Cellular White Matter But Not Extracellular Free Water on Diffusion MRI in Individuals at Clinical High Risk for Psychosis. Am J Psychiatry. 2019;176(10):820–828. doi:10.1176/appi.ajp.2019.18091044

51. Tang Y, Wang J, Zhang T, et al. P300 as an index of transition to psychosis and of remission: Data from a clinical high risk for psychosis study and review of literature. Schizophr Res. 2020;226:74–83. doi:10.1016/j.schres.2019.02.014

52. Wu G, Gan R, Li Z, et al. Real-World Effectiveness and Safety of Antipsychotics in Individuals at Clinical High-Risk for Psychosis: Study Protocol for a Prospective Observational Study (ShangHai at Risk for Psychosis-Phase 2). Neuropsychiatr Dis Treat. 2019;15:3541–3548. doi:10.2147/NDT.S230904

53. Wu G, Tang X, Gan R, et al. Temporal and time–frequency features of auditory oddball response in distinct subtypes of patients at clinical high risk for psychosis. Eur Arch Psychiatry Clin Neurosci. 2022;272(3):449–459. doi:10.1007/s00406-021-01316-1

54. Wu G, Tang X, Gan R, et al. Automatic auditory processing features in distinct subtypes of patients at clinical high risk for psychosis: Forecasting remission with mismatch negativity. Hum Brain Mapp. 2022;43(18):5452–5464. doi:10.1002/hbm.26021

55. Zhang T, Li H, Woodberry KA, et al. Prodromal psychosis detection in a counseling center population in China: An epidemiological and clinical study. Schizophr Res. 2014;152(2-3):391–399. doi:10.1016/j.schres.2013.11.039

56. Zhang T, Li H, Xu L, et al. SU39. Do Baseline Clinical and Neurocognitive Features Predict Conversion in Individuals With Clinical High Risk to Psychosis in Shanghai? Schizophr Bull. 2017;43(suppl_1):S175. doi:10.1093/schbul/sbx024.037

57. Zhang T, Xu L, Tang Y, et al. Prediction of psychosis in prodrome: development and validation of a simple, personalized risk calculator. Psychol Med. 2019;49(12):1990–1998. doi:10.1017/S0033291718002738

58. Zhang T, Xu L, Tang X, et al. Comprehensive review of multidimensional biomarkers in the ShangHai At Risk for Psychosis (SHARP) program for early psychosis identification. Psychiatry Clin Neurosci Rep. 2023;2(4):e152. doi:10.1002/pcn5.152

59. Zhang T, Xu L, Wei Y, et al. Advancements and Future Directions in Prevention Based on Evaluation for Individuals With Clinical High Risk of Psychosis: Insights From the SHARP Study. Schizophr Bull. Published online May 14, 2024:sbae066. doi:10.1093/schbul/sbae066

60. Zhang T, Li H, Tang Y, et al. 21.4 BASELINE CLINICAL AND BIOLOGICAL VARIABLES PREDICTING 1 YEAR OUTCOME OF SUBJECTS AT CLINICAL HIGH RISK OF PSYCHOSIS: INSIGHT FROM SHANGHAI AT RISK FOR PSYCHOSIS (SHARP) PROGRAM. Schizophr Bull. 2018;44(suppl_1):S36. doi:10.1093/schbul/sby014.088

61. Zhang T, Li H, Tang Y, et al. Validating the Predictive Accuracy of the NAPLS-2 Psychosis Risk Calculator in a Clinical High-Risk Sample From the SHARP (Shanghai At Risk for Psychosis) Program. Am J Psychiatry. 2018;175(9):906–908. doi:10.1176/appi.ajp.2018.18010036

62. Zhang T, Raballo A, Zeng J, et al. Antipsychotic prescription, assumption and conversion to psychosis: resolving missing clinical links to optimize prevention through precision. Schizophrenia. 2022;8(1):1–9. doi:10.1038/s41537-022-00254-8

63. Zhang T, Tang X, Li H, et al. Clinical subtypes that predict conversion to psychosis: A canonical correlation analysis study from the ShangHai At Risk for Psychosis program. Aust N Z J Psychiatry. 2020;54(5):482–495. doi:10.1177/0004867419872248

64. Zhang T, Wang J, Xu L, et al. Subtypes of Clinical High Risk for Psychosis that Predict Antipsychotic Effectiveness in Long-Term Remission. Pharmacopsychiatry. 2021;54(1):23–30. doi:10.1055/a-1252-2942

65. Zhang T, Xu L, Chen Y, et al. Conversion to psychosis in adolescents and adults: similar proportions, different predictors. Psychol Med. 2021;51(12):2003–2011. doi:10.1017/S0033291720000756

66. Zhang T, Xu L, Tang X, et al. Real-world effectiveness of antipsychotic treatment in psychosis prevention in a 3-year cohort of 517 individuals at clinical high risk from the SHARP (ShangHai At Risk for Psychosis). Aust N Z J Psychiatry. 2020;54(7):696–706. doi:10.1177/0004867420917449

67. Zhang T, Xu L, Li H, et al. Calculating individualized risk components using a mobile app-based risk calculator for clinical high risk of psychosis: findings from ShangHai At Risk for Psychosis (SHARP) program. Psychol Med. 2021;51(4):653–660. doi:10.1017/S003329171900360X

68. Zhang T, Xu L, Tang Y, et al. Using “WeChat” online social networking in a real-world needs analysis of family members of youths at clinical high risk of psychosis. Aust N Z J Psychiatry. 2018;52(4):375–382. doi:10.1177/0004867417712460

69. Zhang T, Xu L, Tang Y, et al. Duration of untreated prodromal symptoms in a Chinese sample at a high risk for psychosis: demographic, clinical, and outcome. Psychol Med. 2018;48(8):1274–1281. doi:10.1017/S0033291717002707

70. Zhang T, Yang S, Xu L, et al. Poor functional recovery is better predicted than conversion in studies of outcomes of clinical high risk of psychosis: insight from SHARP. Psychol Med. 2020;50(9):1578–1584. doi:10.1017/S0033291719002174

71. Zhang T, Zeng J, Wei Y, et al. Changes in inflammatory balance correlates with conversion to psychosis among individuals at clinical high-risk: A prospective cohort study. Psychiatry Res. 2022;318:114938. doi:10.1016/j.psychres.2022.114938

72. Zhang TH, Li HJ, Woodberry KA, et al. Two-year follow-up of a Chinese sample at clinical high risk for psychosis: timeline of symptoms, help-seeking and conversion. Epidemiol Psychiatr Sci. 2017;26(3):287–298. doi:10.1017/S2045796016000184

73. McGlashan T, Walsh B, Woods S. The Psychosis-Risk Syndrome: Handbook for Diagnosis and Follow-Up. Oxford University Press; 2010.

74. Jones SH, Thornicroft G, Coffey M, Dunn G. A Brief Mental Health Outcome Scale: Reliability and Validity of the Global Assessment of Functioning (GAF). Br J Psychiatry. 1995;166(5):654–659. doi:10.1192/bjp.166.5.654

75. Wechsler D. Wechsler Abbreviated Scale of Intelligence. Published online 1999. doi:10.1037/t15170-000

76. Whitfield-Gabrieli S, Nieto-Castanon A. Conn: A Functional Connectivity Toolbox for Correlated and Anticorrelated Brain Networks. https://home.liebertpub.com/brain. doi:10.1089/brain.2012.0073

77. Jenkinson M, Bannister P, Brady M, Smith S. Improved optimization for the robust and accurate linear registration and motion correction of brain images. NeuroImage. 2002;17(2):825–841. doi:10.1016/s1053-8119(02)91132-8

78. Fox MD, Snyder AZ, Vincent JL, Corbetta M, Van Essen DC, Raichle ME. The human brain is intrinsically organized into dynamic, anticorrelated functional networks. Proc Natl Acad Sci. 2005;102(27):9673–9678. doi:10.1073/pnas.0504136102

79. Benjamini Y, Hochberg Y. Controlling the False Discovery Rate: A Practical and Powerful Approach to Multiple Testing. J R Stat Soc Ser B Methodol. 1995;57(1):289–300.

80. R Core Team. R: A language and environment for statistical computing. Published online 2022. https://www.R-project.org/

81. Wickham H, Averick M, Bryan J, et al. Welcome to the Tidyverse. J Open Source Softw. 2019;4(43):1686. doi:10.21105/joss.01686

82. Kay SR, Sevy S. Pyramidical Model of Schizophrenia. Schizophr Bull. 1990;16(3):537–545.

83. Kim JH, Kim SY, Lee J, Oh KJ, Kim YB, Cho ZH. Evaluation of the factor structure of symptoms in patients with schizophrenia. Psychiatry Res. 2012;197(3):285–289. doi:10.1016/j.psychres.2011.10.006

84. McGorry PD, Bell RC, Dudgeon PL, Jackson HJ. The dimensional structure of first episode psychosis: an exploratory factor analysis. Psychol Med. 1998;28(4):935–947. doi:10.1017/S0033291798006771

85. Ventura J, Nuechterlein KH, Subotnik KL, Gutkind D, Gilbert EA. Symptom dimensions in recent-onset schizophrenia and mania: a principal components analysis of the 24-item Brief Psychiatric Rating Scale. Psychiatry Res. 2000;97(2-3):129–135. doi:10.1016/S0165-1781(00)00228-6

86. Liu H, Kaneko Y, Ouyang X, et al. Schizophrenic Patients and Their Unaffected Siblings Share Increased Resting-State Connectivity in the Task-Negative Network but Not Its Anticorrelated Task-Positive Network. Schizophr Bull. 2012;38(2):285–294. doi:10.1093/schbul/sbq074

87. Hilland E, Johannessen C, Jonassen R, et al. Aberrant default mode connectivity in adolescents with early-onset psychosis: A resting state fMRI study. NeuroImage Clin. 2022;33:102881. doi:10.1016/j.nicl.2021.102881

88. Lee H, Lee DK, Park K, Kim CE, Ryu S. Default mode network connectivity is associated with long-term clinical outcome in patients with schizophrenia. NeuroImage Clin. 2019;22:101805. doi:10.1016/j.nicl.2019.101805

89. Li S, Hu N, Zhang W, et al. Dysconnectivity of Multiple Brain Networks in Schizophrenia: A Meta-Analysis of Resting-State Functional Connectivity. Front Psychiatry. 2019;10. doi:10.3389/fpsyt.2019.00482

90. Chan SY, Brady R, Hwang M, et al. Heterogeneity of Outcomes and Network Connectivity in Early-Stage Psychosis: A Longitudinal Study. Schizophr Bull. 2020;47(1):138–148. doi:10.1093/schbul/sbaa079

91. Kujawa A, Burkhouse KL, Karich SR, Fitzgerald KD, Monk CS, Phan KL. Reduced Reward Responsiveness Predicts Change in Depressive Symptoms in Anxious Children and Adolescents Following Treatment. J Child Adolesc Psychopharmacol. 2019;29(5):378–385. doi:10.1089/cap.2018.0172

92. Scholing A, Emmelkamp PMG. Prediction of treatment outcome in social phobia: a cross-validation. Behav Res Ther. 1999;37(7):659–670. doi:10.1016/S0005-7967(98)00175-2

93. Whitfield-Gabrieli S, Ghosh SS, Nieto-Castanon A, et al. Brain connectomics predict response to treatment in social anxiety disorder. Mol Psychiatry. 2016;21(5):680–685. doi:10.1038/mp.2015.109

94. Berman MG, Peltier S, Nee DE, Kross E, Deldin PJ, Jonides J. Depression, rumination and the default network. Soc Cogn Affect Neurosci. 2011;6(5):548–555. doi:10.1093/scan/nsq080

95. Hamilton JP, Farmer M, Fogelman P, Gotlib IH. Depressive Rumination, the Default-Mode Network, and the Dark Matter of Clinical Neuroscience. Biol Psychiatry. 2015;78(4):224–230. doi:10.1016/j.biopsych.2015.02.020

96. Jacobs RH, Jenkins LM, Gabriel LB, et al. Increased Coupling of Intrinsic Networks in Remitted Depressed Youth Predicts Rumination and Cognitive Control. PLOS ONE. 2014;9(8):e104366. doi:10.1371/journal.pone.0104366

97. Posner J, Cha J, Wang Z, et al. Increased Default Mode Network Connectivity in Individuals at High Familial Risk for Depression. Neuropsychopharmacology. 2016;41(7):1759–1767. doi:10.1038/npp.2015.342

98. Sheline YI, Price JL, Yan Z, Mintun MA. Resting-state functional MRI in depression unmasks increased connectivity between networks via the dorsal nexus. Proc Natl Acad Sci. 2010;107(24):11020–11025. doi:10.1073/pnas.1000446107

99. Kucyi A, Moayedi M, Weissman-Fogel I, et al. Enhanced Medial Prefrontal-Default Mode Network Functional Connectivity in Chronic Pain and Its Association with Pain Rumination. J Neurosci. 2014;34(11):3969–3975. doi:10.1523/JNEUROSCI.5055-13.2014

100. Lynch CJ, Uddin LQ, Supekar K, Khouzam A, Phillips J, Menon V. Default Mode Network in Childhood Autism: Posteromedial Cortex Heterogeneity and Relationship with Social Deficits. Biol Psychiatry. 2013;74(3):212–219. doi:10.1016/j.biopsych.2012.12.013

101. Patriat R, Birn RM, Keding TJ, Herringa RJ. Default-Mode Network Abnormalities in Pediatric Posttraumatic Stress Disorder. J Am Acad Child Adolesc Psychiatry. 2016;55(4):319–327. doi:10.1016/j.jaac.2016.01.010

102. Zhou HX, Chen X, Shen YQ, et al. Rumination and the default mode network: Meta-analysis of brain imaging studies and implications for depression. NeuroImage. 2020;206:116287. doi:10.1016/j.neuroimage.2019.116287

103. Chai XJ, Hirshfeld-Becker D, Biederman J, et al. Altered Intrinsic Functional Brain Architecture in Children at Familial Risk of Major Depression. Biol Psychiatry. 2016;80(11):849–858. doi:10.1016/j.biopsych.2015.12.003

104. Li BJ, Friston K, Mody M, Wang HN, Lu HB, Hu DW. A brain network model for depression: From symptom understanding to disease intervention. CNS Neurosci Ther. 2018;24(11):1004–1019. doi:10.1111/cns.12998

105. Zhu X, Zhu Q, Shen H, Liao W, Yuan F. Rumination and Default Mode Network Subsystems Connectivity in First-episode, Drug-Naive Young Patients with Major Depressive Disorder. Sci Rep. 2017;7(1):43105. doi:10.1038/srep43105

106. Watkins ER, Roberts H. Reflecting on rumination: Consequences, causes, mechanisms and treatment of rumination. Behav Res Ther. 2020;127:103573. doi:10.1016/j.brat.2020.103573

107. Ehring T, Watkins ER. Repetitive Negative Thinking as a Transdiagnostic Process. Int J Cogn Ther. 2008;1(3):192–205. doi:10.1521/ijct.2008.1.3.192

108. Addington J, Cornblatt BA, Cadenhead KS, et al. At Clinical High Risk for Psychosis: Outcome for Nonconverters. Am J Psychiatry. 2011;168(8):800–805. doi:10.1176/appi.ajp.2011.10081191

109. Beck K, Andreou C, Studerus E, et al. Clinical and functional long-term outcome of patients at clinical high risk (CHR) for psychosis without transition to psychosis: A systematic review. Schizophr Res. 2019;210:39–47. doi:10.1016/j.schres.2018.12.047

110. McFadden KL, Cornier MA, Melanson EL, Bechtell JL, Tregellas JR. Effects of exercise on resting-state default mode and salience network activity in overweight/obese adults. Neuroreport. 2013;24(15):866–871. doi:10.1097/WNR.0000000000000013

111. Brewer JA, Worhunsky PD, Gray JR, Tang YY, Weber J, Kober H. Meditation experience is associated with differences in default mode network activity and connectivity. Proc Natl Acad Sci. 2011;108(50):20254–20259. doi:10.1073/pnas.1112029108

112. Fang A, Baran B, Feusner JD, et al. Self-Focused Brain Predictors of Cognitive Behavioral Therapy Response in a Transdiagnostic Sample. medRxiv. Published online September 2, 2023:2023.08.30.23294878. doi:10.1101/2023.08.30.23294878

113. Posner J, Hellerstein DJ, Gat I, et al. Antidepressants Normalize the Default Mode Network in Patients With Dysthymia. JAMA Psychiatry. 2013;70(4):373–382. doi:10.1001/jamapsychiatry.2013.455

114. Liston C, Chen AC, Zebley BD, et al. Default Mode Network Mechanisms of Transcranial Magnetic Stimulation in Depression. Biol Psychiatry. 2014;76(7):517–526. doi:10.1016/j.biopsych.2014.01.023

115. Zhang J, Raya J, Morfini F, et al. Reducing default mode network connectivity with mindfulness-based fMRI neurofeedback: a pilot study among adolescents with affective disorder history. Mol Psychiatry. Published online March 30, 2023:1–9. doi:10.1038/s41380-023-02032-z

116. Fioravanti M, Bianchi V, Cinti ME. Cognitive deficits in schizophrenia: an updated metanalysis of the scientific evidence. BMC Psychiatry. 2012;12(1):64. doi:10.1186/1471-244X-12-64

117. Seidman LJ. Schizophrenia and Brain Dysfunction: An Integration of Recent Neurodiagnostic Findings. Psychol Bull. 1983;94(2):195–238.

118. Cella M, Preti A, Edwards C, Dow T, Wykes T. Cognitive remediation for negative symptoms of schizophrenia: A network meta-analysis. Clin Psychol Rev. 2017;52:43–51. doi:10.1016/j.cpr.2016.11.009

119. Harvey PD, Koren D, Reichenberg A, Bowie CR. Negative Symptoms and Cognitive Deficits: What Is the Nature of Their Relationship? Schizophr Bull. 2006;32(2):250–258. doi:10.1093/schbul/sbj011

120. Lin CH, Huang CL, Chang YC, et al. Clinical symptoms, mainly negative symptoms, mediate the influence of neurocognition and social cognition on functional outcome of schizophrenia. Schizophr Res. 2013;146(1):231–237. doi:10.1016/j.schres.2013.02.009

121. Thomas ML, Green MF, Hellemann G, et al. Modeling Deficits From Early Auditory Information Processing to Psychosocial Functioning in Schizophrenia. JAMA Psychiatry. 2017;74(1):37–46. doi:10.1001/jamapsychiatry.2016.2980

122. Andrews-Hanna JR, Reidler JS, Sepulcre J, Poulin R, Buckner RL. Functional-Anatomic Fractionation of the Brain’s Default Network. Neuron. 2010;65(4):550–562. doi:10.1016/j.neuron.2010.02.005

123. Whitfield-Gabrieli S, Ford JM. Default Mode Network Activity and Connectivity in Psychopathology. Annu Rev Clin Psychol. 2012;8(1):49–76. doi:10.1146/annurev-clinpsy-032511-143049

124. Shenton ME, Dickey CC, Frumin M, McCarley RW. A review of MRI findings in schizophrenia. Schizophr Res. 2001;49(1-2):1–52.

125. Ford JM, Roach BJ, Jorgensen KW, et al. Tuning in to the Voices: A Multisite fMRI Study of Auditory Hallucinations. Schizophr Bull. 2009;35(1):58–66. doi:10.1093/schbul/sbn140

126. Bauer CCC, Okano K, Gosh SS, et al. Real-time fMRI neurofeedback reduces auditory hallucinations and modulates resting state connectivity of involved brain regions: Part 2: Default Mode Network -Preliminary evidence-. Psychiatry Res. 2020;284:112770. doi:10.1016/j.psychres.2020.112770

127. Hafner H, Maurer K, Loffler W, Der Heiden WA, Hambrecht M, Schultze-Lutter F. Modeling the Early Course of Schizophrenia. Schizophr Bull. 2003;29(2):325–340. doi:10.1093/oxfordjournals.schbul.a007008

128. Iyer SN, Boekestyn L, Cassidy CM, King S, Joober R, Malla AK. Signs and symptoms in the pre-psychotic phase: description and implications for diagnostic trajectories. Psychol Med. 2008;38(8):1147–1156. doi:10.1017/S0033291708003152

