## Supplementary Materials for "Dissociable Default Mode Network Connectivity Patterns Underlie Distinct Symptoms in Psychosis Risk"

| Chelsea Ajunwa | Jiahe Zhang |
|---|---|
| Northeastern University | Northeastern University |
| 360 Huntington Ave, 129 ISEC | 360 Huntington Ave, 129 ISEC |
| Boston, MA 02120 | Boston, MA 02120 |
| | |

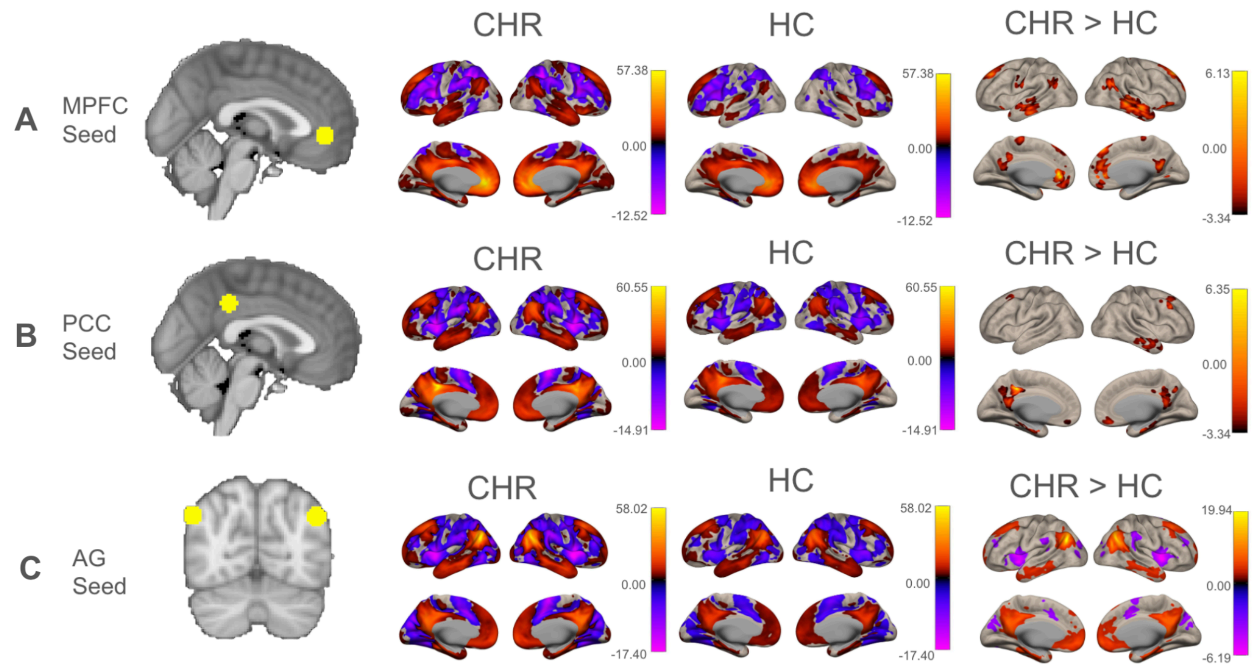


**Supplementary Figure 1.** Functional connectivity of key DMN nodes in clinical high-risk (CHR) vs healthy controls (HC). **A)** An 8mm sphere in the medial prefrontal cortex (MPFC) was used as the seed in a whole-brain seed-to-voxel analysis. This analysis revealed greater connectivity between the MPFC and several regions, including the superior temporal gyrus (STG), middle temporal gyrus (MTG), and posterior cingulate cortex (PCC) ($p$ <.001, uncorrected) in CHR compared to controls. **B)** An 8mm sphere in the PCC was used as the seed in a whole-brain seed-to-voxel analysis. CHR exhibited greater connectivity between the PCC and regions such as the MPFC. C) Two 8mm spheres in the bilateral angular gyri (AG) were used as the seed in a whole-brain seed-to-voxel analysis. CHR exhibited greater connectivity between the AG and DMN regions such as the MPFC and PCC.

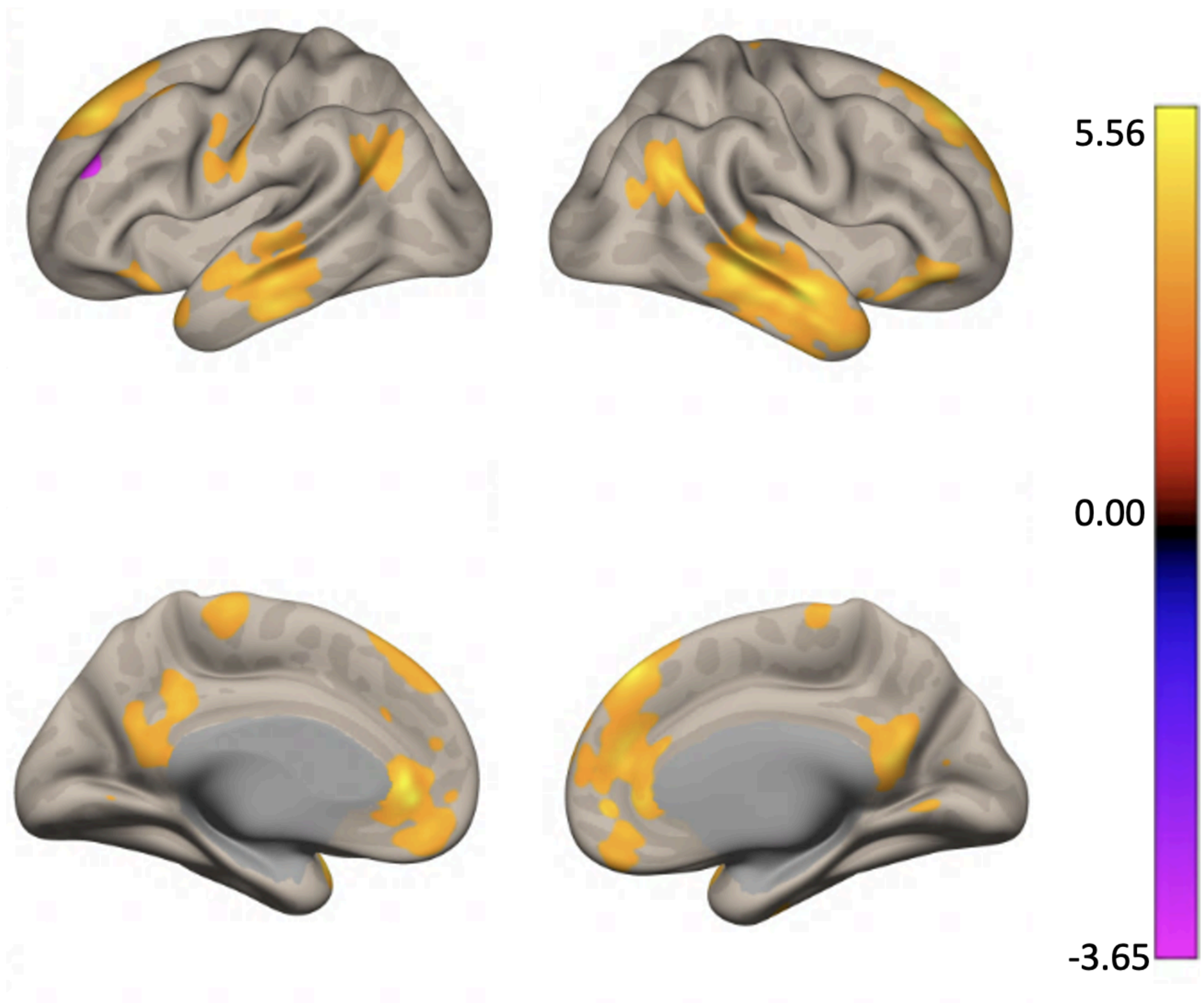


**Supplementary Figure 2.** MPFC hyperconnectivity in CHR after accounting for age, sex, and average framewise displacement covariates. A whole brain search again revealed that, compared to HC, CHR exhibited elevated connectivity between the MPFC seed and several regions. This includes bilateral auditory cortices (middle temporal gyrus and superior temporal gyrus) and the posterior cingulate cortex (*p* <.001, uncorrected). Other smaller clusters also exhibited hyperconnectivity to the MPFC seed, including additional DMN regions such as bilateral angular gyri, superior frontal gyri and left cerebellum Crus II.

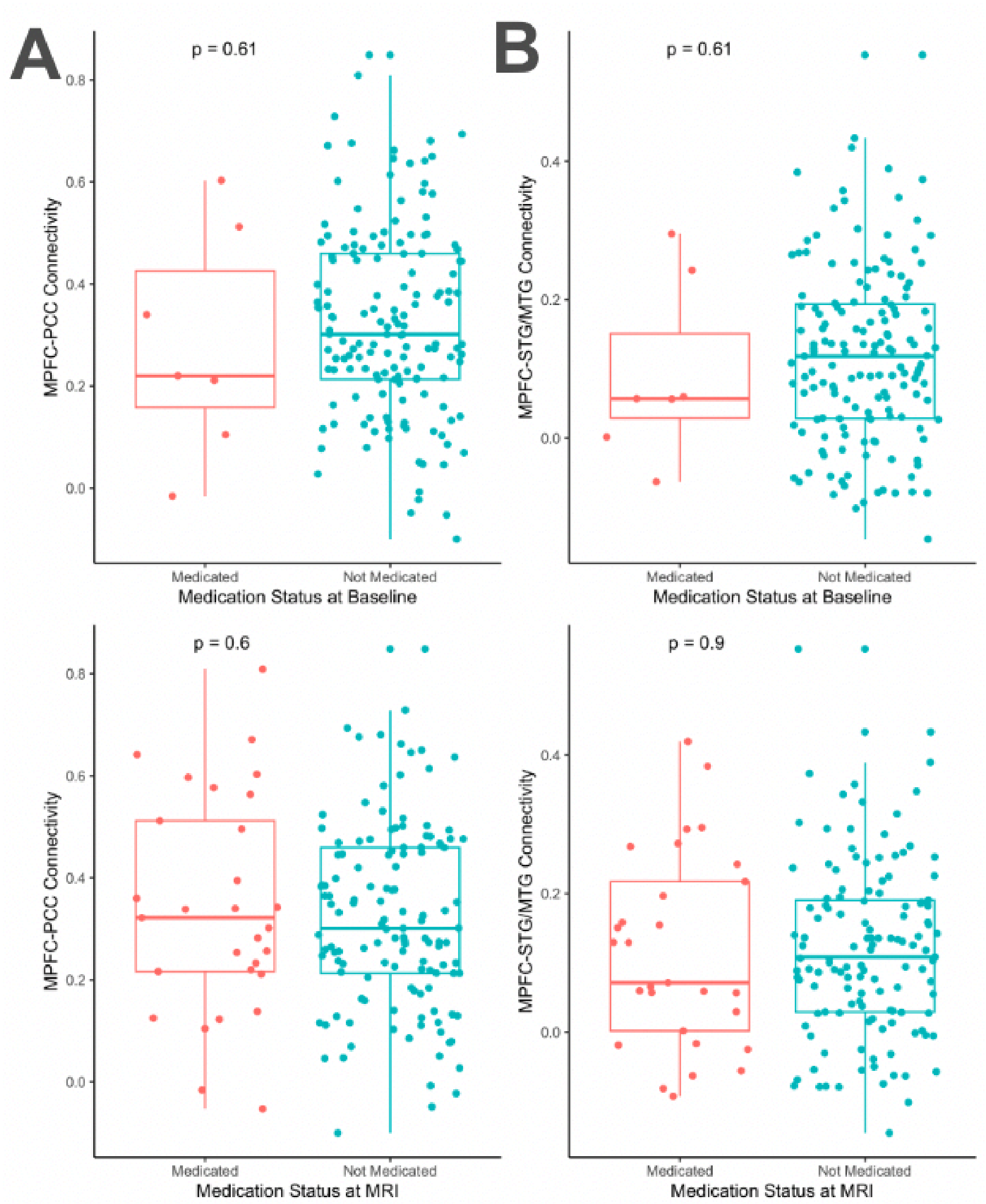


**Supplementary Figure 3.** MPFC-PCC and MPFC-STG/MTG connectivity in medicated vs non-medicated subjects. **A)** MPFC-PCC connectivity in medicated subjects (n=7 at baseline, n=29 at MRI) is not significantly different from MPFC-PCC connectivity in non-medicated subjects (n=151 at baseline, n=129 at MRI ). **B)** MPFC-STG/MTG connectivity in medicated subjects (n=7 at baseline, n=29 at MRI) is not significantly different from MPFC-STG/MTG connectivity in non-medicated subjects (n=151 at baseline, n=129 at MRI ).

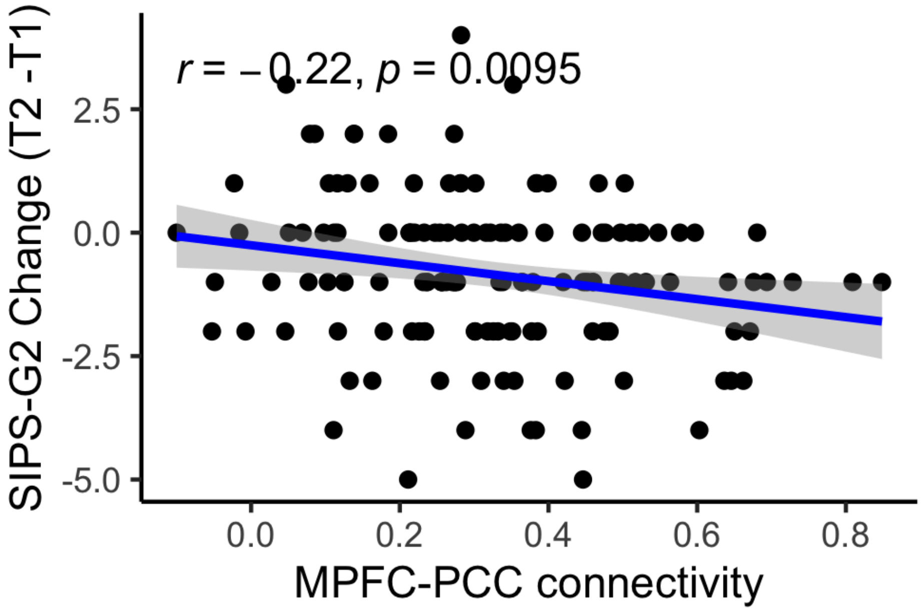


**Supplementary Figure 4.** Relationship between MPFC-PCC connectivity and change in SIPS-G2 (dysphoric mood). MPFC-PCC connectivity was derived from CHR>HC comparison, independently of symptom measures. In CHR, individual differences in MPFC-PCC connectivity were significantly associated with variation in SIPS-G2 change. Change in SIPS-G2 was calculated based on SIPS-G2 at follow-up (one year post-baseline) and SIPS-G2 at baseline assessment.

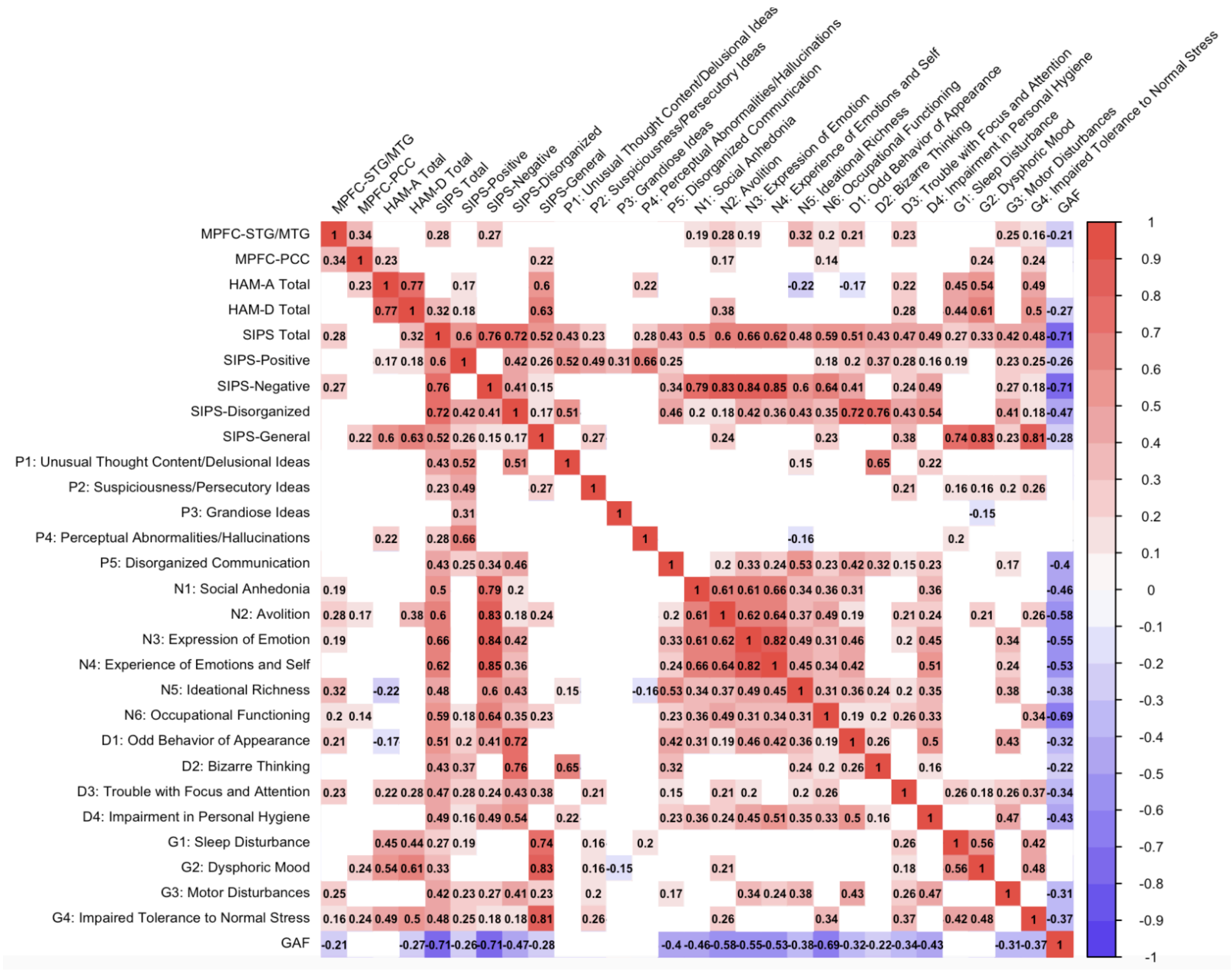


**Supplementary Figure 5**. Correlation matrix with relevant SIPS items. Pearson's correlation coefficients are displayed. Positive correlations are displayed in red and negative correlations are displayed in blue. Non-significant correlations are blank (*p* < 0.5 uncorrected). Visualization was created using the *corrplot* package in R v4.2.2.(Wei & Simko, 2021)

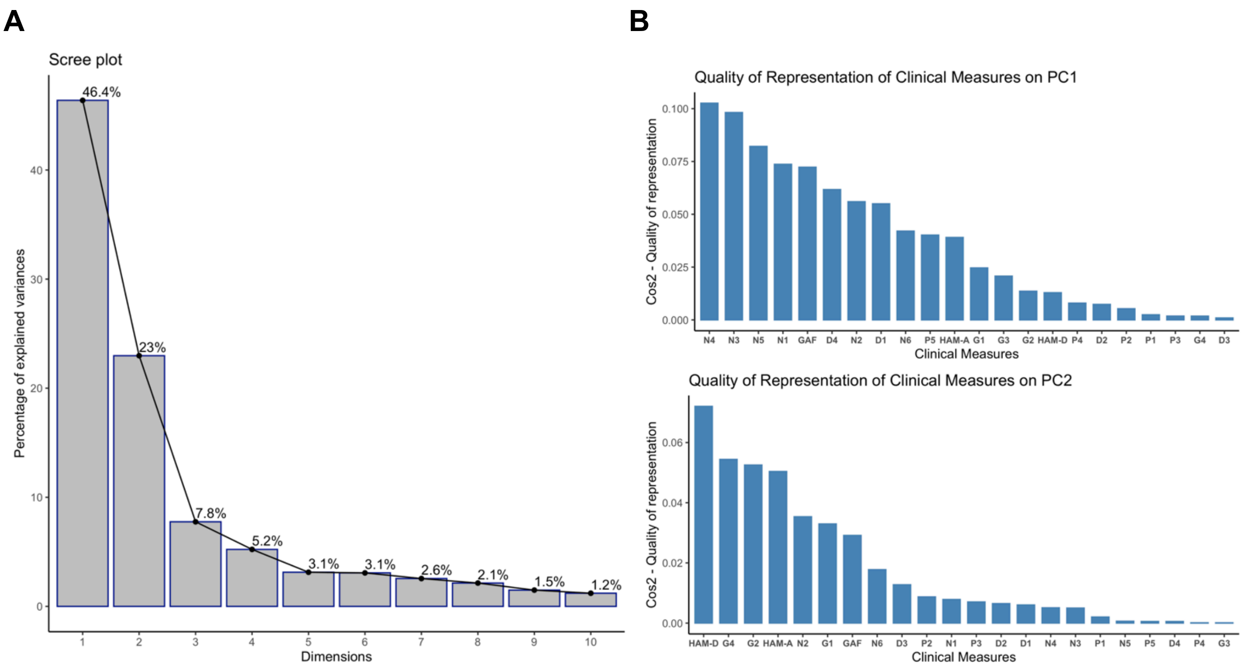


**Supplementary Figure 6**. Principal components derived from symptom data. **A)** Scree plot indicates the amount of variance explained by each principal component. The first principal component explains 46.4% of the variance in the symptom data. The second principal component explains 23% of the variance in the symptom data. **B)** Plot indicates quality of representation of clinical measures (including HAM-A, HAM-D, GAF, and the 19 SIPS items) on the first two principal components. Quality of representation (cos2) is indicated on the y-axis. Visualizations were created using the *factoextra* package in R v4.2.2.(Alboukadel Kassambara & Mundt, 2020)

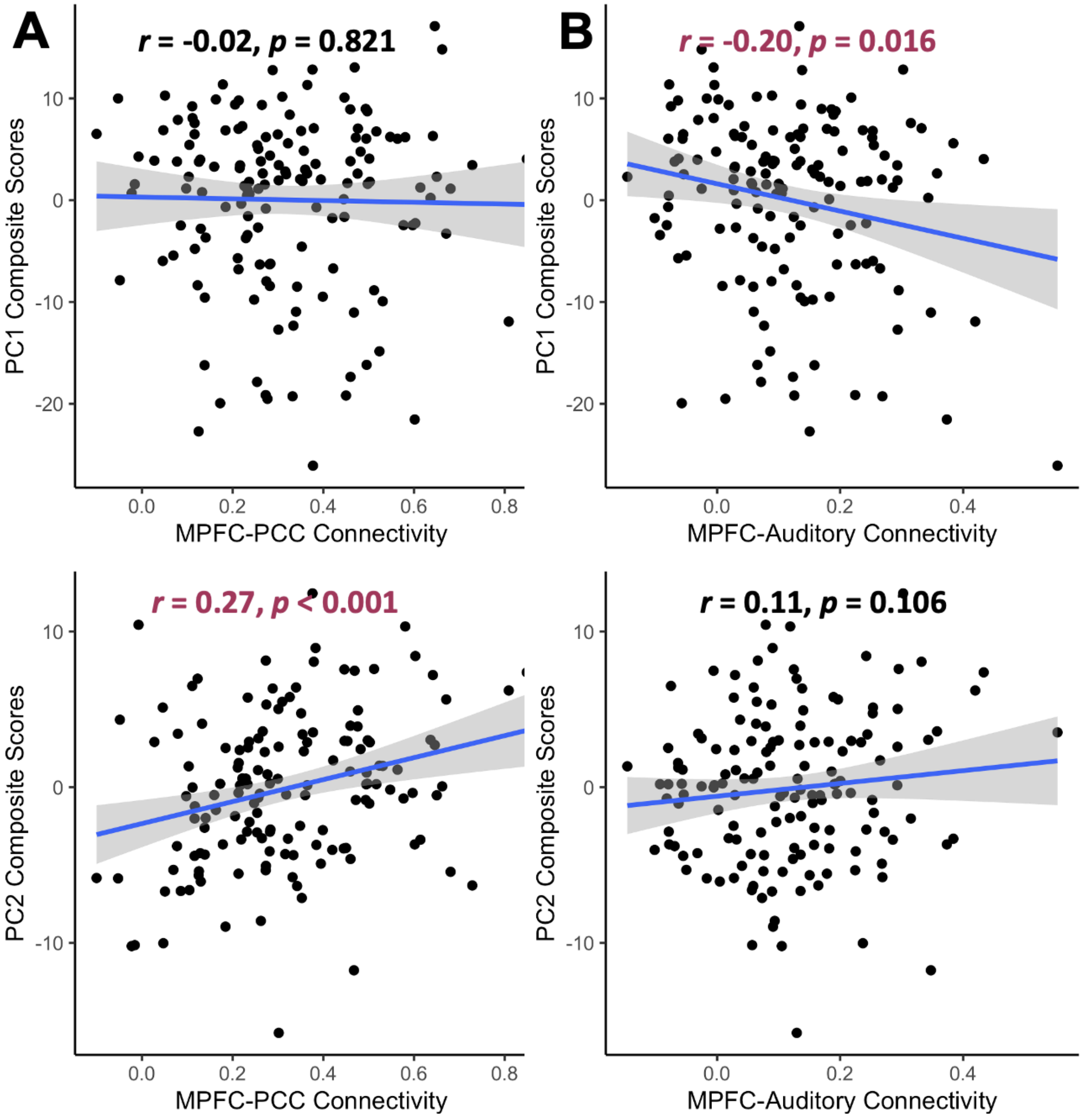


**Supplementary Figure 7**. Dissociable relationships between MPFC connectivity and symptom dimensions in CHR. **A)** Individual differences in MPFC-PCC connectivity were significantly associated with variation in PC2 composite scores, but not in PC1 scores. **B)** Individual differences in MPFC-STG/MTG connectivity were associated with variation in PC1 composite scores, but not in PC2 composite scores.

**Supplementary Table 1. Significant clusters exhibiting higher functional connectivity to the MPFC seed in CHR vs. HC.**

| | Peak Location | Location | MNI Coordinates (x y z) | Size | Size p-FWE | Size p-FDR | Size p-unc | Peak p-FWE | Peak p-unc |
|---|---|---|---|---|---|---|---|---|---|
| 1 | Right Medial Temporal Gyrus | Right Auditory Cortex | (56 -6 -22) | 333 4 | 0.000 | 0.000 | 0.000 | 0.002 | 0.000 |
| 2 | Dorsal Anterior Cingulate Cortex | Medial Prefrontal Cortex | (4 42 0) | 301 0 | 0.000 | 0.000 | 0.000 | 0.000 | 0.000 |
| 3 | Left Medial Temporal Gyrus | Left Auditory Cortex | (-52 -24 -20) | 1165 | 0.000 | 0.000 | 0.000 | 0.080 | 0.000 |
| 4 | Posterior Cingulate Cortex | Posterior Cingulate Cortex | (2 -58 06) | 677 | 0.000 | 0.000 | 0.000 | 0.132 | 0.000 |
| 5 | Right Angular Gyrus | Right Angular Gyrus | (42 -50 24) | 563 | 0.000 | 0.000 | 0.000 | 0.182 | 0.000 |
| 6 | Left Angular Gyrus | Left Angular Gyrus | (-44 -62 30) | 361 | 0.000 | 0.000 | 0.000 | 0.823 | 0.000 |
| 7 | Left and Right Precentral Gyrus | Primary Motor Cortex | (-6 -22 66) | 206 | 0.000 | 0.002 | 0.000 | 0.482 | 0.000 |
| 8 | Left Temporal Pole | Left Temporal Pole | (-40 16 -22) | 156 | 0.001 | 0.001 | 0.000 | 0.188 | 0.000 |
| 9 | Left Cerebellum Crus II | Left Cerebellum Crus II | (-28 -88 -40) | 140 | 0.012 | 0.002 | 0.000 | 0.335 | 0.000 |
| 10 | Left Precentral Gyrus | Left Precentral and Postcentral Gyri | (-66 -2 20) | 136 | 0.010 | 0.002 | 0.000 | 0.445 | 0.000 |
| 11 | Middle Frontal Gyrus | Middle Frontal Gyrus | (-32 20 50) | 68 | 0.18 | 0.041 | 0.005 | 0.999 | 0.000 |

**Supplementary Table 2. Sensitivity analysis results.**

| Outcome variables | β | Uncorrected p-value | Uncorrected p-value, controlling for age, sex, and framewise displacement | $q_{FDR}$-value, controlling for age, sex, and framewise displacement |
|---|---|---|---|---|
| **MPFC-PCC connectivity as predictor** | | | | |
| **HAM-A Total** | **5.213** | **0.006** | **0.004** | **0.048** |
| HAM-D Total | 5.949 | 0.057 | 0.058 | 0.171 |
| P1: Unusual Thought Content/Delusional Ideas | -0.264 | 0.716 | 0.729 | 0.937 |
| P2: Suspiciousness/Persecutory Ideas | 0.031 | 0.966 | 0.966 | 0.983 |
| P3: Grandiose Ideas | -0.563 | 0.085 | 0.084 | 0.212 |
| P4: Perceptual Abnormalities/Hallucinations | -0.488 | 0.627 | 0.555 | 0.823 |
| P5: Disorganized Communication | 0.086 | 0.845 | 0.836 | 0.937 |
| N1: Social Anhedonia | 1.147 | 0.051 | 0.045 | 0.154 |
| N2: Avolition | 1.312 | 0.022 | 0.017 | 0.091 |
| N3: Expression of Emotion | 1.123 | 0.081 | 0.064 | 0.171 |
| N4: Experience of Emotions and Self | -0.083 | 0.873 | 0.879 | 0.937 |
| N5: Ideational Richness | 0.070 | 0.867 | 0.849 | 0.937 |
| N6: Occupational Functioning | 1.466 | 0.033 | 0.031 | 0.124 |
| D1: Odd Behavior of Appearance | 0.145 | 0.786 | 0.785 | 0.937 |
| D2: Bizarre Thinking | 0.261 | 0.773 | 0.746 | 0.937 |
| D3: Trouble with Focus and Attention | -0.074 | 0.879 | 0.859 | 0.937 |
| D4: Impairment in Personal Hygiene | 0.072 | 0.847 | 0.823 | 0.937 |
| G1: Sleep Disturbance | 0.544 | 0.265 | 0.257 | 0.457 |
| **G2: Dysphoric Mood** | **1.983** | **0.001** | **<0.001** | **<0.001** |
| G3: Motor Disturbances | -0.287 | 0.257 | 0.250 | 0.457 |
| **G4: Impaired Tolerance to Normal Stress** | **1.909** | **0.001** | **0.001** | **0.016** |
| SIPS-N | 5.034 | 0.056 | 0.045 | 0.154 |

| | | | | |
|---|---|---|---|---|
| SIPS-P | -1.198 | 0.448 | 0.434 | 0.672 |
| GAF | -6.643 | 0.062 | 0.058 | 0.171 |
| **MPFC-STG/MTG connectivity as predictor** | | | | |
| HAM-A Total | 2.682 | 0.286 | 0.342 | 0.581 |
| HAM-D Total | 1.442 | 0.552 | 0.764 | 0.937 |
| P1: Unusual Thought Content/Delusional Ideas | 0.603 | 0.780 | 0.600 | 0.847 |
| P2: Suspiciousness/Persecutory Ideas | 1.352 | 0.275 | 0.213 | 0.426 |
| P3: Grandiose Ideas | 0.204 | 0.437 | 0.679 | 0.905 |
| P4: Perceptual Abnormalities/Hallucinations | 0.591 | 0.374 | 0.636 | 0.872 |
| P5: Disorganized Communication | 0.742 | 0.164 | 0.233 | 0.447 |
| N1: Social Anhedonia | 1.464 | 0.027 | 0.091 | 0.218 |
| **N2: Avolition** | **2.307** | **0.001** | **0.005** | **0.048** |
| N3: Expression of Emotion | 1.239 | 0.047 | 0.177 | 0.369 |
| N4: Experience of Emotions and Self | 0.650 | 0.215 | 0.431 | 0.672 |
| **N5: Ideational Richness** | **2.028** | **<0.001** | **<0.001** | **<0.001** |
| N6: Occupational Functioning | 2.584 | 0.008 | 0.011 | 0.066 |
| D1: Odd Behavior of Appearance | 1.326 | 0.019 | 0.096 | 0.219 |
| D2: Bizarre Thinking | 0.111 | 0.854 | 0.927 | 0.967 |
| D3: Trouble with Focus and Attention | 1.157 | 0.011 | 0.063 | 0.171 |
| D4: Impairment in Personal Hygiene | 0.454 | 0.317 | 0.351 | 0.581 |
| G1: Sleep Disturbance | 0.416 | 0.817 | 0.566 | 0.823 |
| G2: Dysphoric Mood | 0.019 | 0.785 | 0.983 | 0.983 |
| G3: Motor Disturbances | 0.963 | 0.003 | 0.010 | 0.066 |
| G4: Impaired Tolerance to Normal Stress | 2.053 | 0.039 | 0.019 | 0.091 |
| **SIPS-N** | **10.272** | **0.001** | **0.006** | **0.048** |
| SIPS-P | 3.491 | 0.086 | 0.130 | 0.283 |
| GAF | 11.524 | 0.010 | 0.029 | 0.124 |

**Supplementary Table 3. Principal component analysis variable loadings.**

| Variables | PC1 Loadings | PC2 Loadings |
|---|---|---|
| HAM-A Total | 0.22 | 0.35 |
| HAM-D Total | 0.13 | 0.42 |
| P1: Unusual Thought Content/Delusional Ideas | -0.05 | -0.07 |
| P2: Suspiciousness/Persecutory Ideas | 0.08 | 0.15 |
| P3: Grandiose Ideas | 0.05 | -0.13 |
| P4: Perceptual Abnormalities/Hallucinations | 0.10 | 0.01 |
| P5: Disorganized Communication | -0.22 | -0.04 |
| N1: Social Anhedonia | -0.30 | 0.14 |
| N2: Avolition | -0.26 | 0.29 |
| N3: Expression of Emotion | -0.35 | 0.11 |
| N4: Experience of Emotions and Self | -0.35 | 0.11 |
| N5: Ideational Richness | -0.32 | -0.04 |
| N6: Occupational Functioning | -0.23 | 0.21 |
| D1: Odd Behavior of Appearance | -0.26 | -0.12 |
| D2: Bizarre Thinking | -0.09 | -0.13 |
| D3: Trouble with Focus and Attention | -0.03 | 0.18 |
| D4: Impairment in Personal Hygiene | -0.27 | -0.04 |
| G1: Sleep Disturbance | 0.17 | 0.28 |
| G2: Dysphoric Mood | 0.13 | 0.36 |
| G3: Motor Disturbances | -0.16 | -0.01 |
| G4: Impaired Tolerance to Normal Stress | 0.05 | 0.37 |
| GAF | 0.30 | -0.27 |

**References**

Alboukadel Kassambara, & Mundt, F. (2020). _*factoextra: Extract and Visualize the Results of Multivariate Data Analyses*_. (R package version 1.0.7) [Computer software]. https://CRAN.R-project.org/package=factoextra

Wei, T., & Simko, V. (2021). *R package “corrplot”: Visualization of a Correlation Matrix* (Version 0.92) [Computer software]. https://github.com/taiyun/corrplot
